# A quantitative analysis of objective feather colour assessment: measurements in the lab are more reliable than in the field

**DOI:** 10.1101/007914

**Authors:** Iker Vaquero-Alba, Andrew McGowan, Daniel Pincheira-Donoso, Matthew R. Evans, Sasha R.X. Dall

## Abstract

The evolution of animal colouration is importantly driven by sexual selection operating on traits used to transmit information to rivals and potential mates, which therefore, have major impacts on fitness. Reflectance spectrometry has become a standard colour-measuring tool, especially after the discovery of tetrachromacy in birds and their ability to detect UV. Birds’ plumage patterns may be invisible to humans, necessitating a reliable and objective way of assessing colouration not dependent on human vision. Plumage colouration measurements can be taken directly on live birds in the field or in the lab (e.g. on collected feathers). Therefore, it is essential to determine which sampling method yields more repeatable and reliable measures, and which of the available quantitative approaches best assess the repeatability of these measures. Using a spectrophotometer, we measured melanin-based colouration in barn swallows’ (*Hirundo rustica*) plumage. We assessed the repeatability of measures obtained with both traditional sampling methods separately to quantitatively determine their reliability. We used the ANOVA-based method for calculating the repeatability of measurements from two years separately, and the GLMM-based method to calculate overall adjusted repeatabilities for both years. We repeated the assessment for the whole reflectance spectrum range and only the human-visible part, to assess the influence of the UV component on the reliabilities of sampling methodologies. Our results reveal very high repeatability for lab measurements and a lower, still moderate to high repeatability, for field measurements. Both increased when limited to only the human-visible part, for all plumage patches except the throat, where we observed the opposite trend. Repeatability between sampling methods was quite low including the whole spectrum, but moderate including only the human-visible part. Our results suggest higher reliability for measurements in the lab and higher power and accuracy of the GLMM-based method. They also suggest UV reflectance differences amongst different plumage patches.

## Introduction

Colour vision involves the capacity to discriminate amongst different wavelengths of light, independent of their intensity [1,2]. Although colouration traits expressed in animals have proven essential components to understand the nature of selection, sexual selection in particular, only relatively recently have scientists appreciated the importance of a systematic understanding of both function and evolution of colouration, as well as the mechanisms that underpin it [3]. Birds in particular, due to their colourful displays and the role of their colour signals in fitness differentials, have traditionally been employed as prime model systems to understand the causes and implications of colour evolution. However, the mechanisms of colour vision and spectral information processing needed to understand how birds perceive colours remain areas with more questions than answers.

Mate choice theory predicts that elaborately ornamented males can provide female birds with direct (if ornamental traits reflect individual condition, useful individual attributes or somatic quality independent of condition) and/or indirect fitness benefits (‘good genes’ or attractiveness for offspring – as conspicuous and costly male traits indicate highly heritable viability) [4–6]. Therefore, birds with more elaborate colourful displays are expected to enjoy a selective advantage given their higher mating chances [4,7].

Given the paramount significance of studies of bird plumage colouration in behavioural and evolutionary ecology, methods for reliably and objectively quantifying such colouration are critical. Methods traditionally used for colour assessment include colour ranks on an arbitrary scale [8], tristimulus colour models based on human vision, such as the Munsell system [9,10], reference colour swatches [11], or digital photography [12–14]. However, although all these methods offer simple and affordable ways of colour measurements in different analytical settings, they lack reliability and objectivity [15], as they are tuned to the human visual system instead of the bird visual system.

Birds do not perceive colours in the same way as humans [16]. Birds have evolved a fourth cone type in their eyes, with a pigment that is sensitive to ultraviolet light. And although we are still far from understanding exactly how colours are perceived by birds [3], progress is being made towards understanding how colour vision works in general and how spectral information is processed by birds and other non-human animals [2,17,18]. Methods have been developed for comparing colour patterns as birds see them, using known properties of bird eyes and generating detailed formal descriptions of colour spaces and the equations used to plot them [19].

Since the 1990s, a wide range of further methods for analyzing spectrophotometry data have emerged [3]. This development stems largely from the revival of interest in UV vision and tetrachromacy in birds and the fact that birds can see colours that humans cannot experience [20,21]. This raises the possibility of the existence of plumage patterns invisible to the human eye, and mate choice behaviours based on the ultraviolet part of the bird reflectance spectrum have been discovered [22,23]. Therefore, reliable and objective ways of quantifying bird coloration, not dependent on human vision, are at a premium. Miniature diode-array spectroradiometer systems, lighter, more portable and affordable than previous spectrometry systems but as precise and objective in colour quantification, have provided popular tools for colour communication studies [3].

Two traditional ways of assessing bird plumage colouration with spectrophotometers have been reported in the literature. Measurements may be taken directly on the bird, applying the pointer of the spectrophotometer to plumage patches as they occur *in situ* [24–29]. Alternatively, measurements may be taken in the lab, with feather samples collected from the field, applying the pointer (the cone-shaped piece of black plastic on top of the probe used to direct the light from the source in a given angle) to “plumage patches” created by mounting these feathers on a flat surface in a way that mimics the original plumage structure [30–38]. Despite the popularity of the use of spectrophotometers for colour assessment and the growing number of studies on bird colouration, few studies have rigorously assessed the consistency of both methods for measuring the colouration of plumage patches, and the repeatability of results obtained when using either one or the other (see [39] for a comparison of carotenoid-based plumage coloration in great tits).

Additionally, there is little consensus on how to best quantify the reliability, or repeatability, of spectral measurements. The most common measure of repeatability, or more precisely, the coefficient of intraclass correlation (*r*_i_), can be formally defined as the proportion of the total variance explained by differences among groups [40,41]:

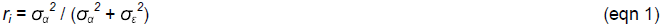

where *σ_α_^2^* is the between-group variance and *σ_ε_^2^* the within-group variance, whereas the sum of both comprises the total phenotypic variance [41]. Until recently, the most common ways to estimate repeatabilities from data with Gaussian errors have employed the correlation-based method [40] or the ANOVA-based method, commonly used by behavioural and evolutionary ecologists [42,43]. However, Nakagawa and Schielzeth [41] developed an innovative R-based function for calculating GLMM-based repeatability estimates, which allows for confounding variables to be factored out and calculates the confidence intervals for each repeatability calculation, inferred from distributions of repeatabilities obtained by parametric bootstrapping.

Our aim is twofold. We first compare two different methods for measuring melanin-based plumage ornamentation to determine which one allows to obtain more repeatable and reliable measures. The methods consist in measuring the feather colouration either directly on the bird in the field or on feather samples in the laboratory. We then compare two statistical methods to assess repeatability, one ANOVA-based and another one GLMM-based, and determine the pros and cons of each of them. We hypothesize that measuring feather colouration in the lab will yield more repeatable and reliable measures, as it avoids the logistic, technical and animal welfare limitations imposed by the field method and provides more controlled conditions during the measurements. Also, we predict more realistic and accurate repeatability estimates with the GLMM-based statistical method, as it allows us to introduce more sources of variation in each analysis. We use the European barn swallow *Hirundo rustica* as a model species. To the best of our knowledge, this is the first time GLMM-based repeatability estimates have been used to assess the reliability of melanin-based plumage colouration measurements.

## Methods

### Field work and data collection

Field work was carried out during May-August 2009 and March-August 2010 in multiple sites, mostly farmlands, around the Falmouth area in Cornwall, UK. A total of 59 adult European barn swallows (21 in 2009 and 38 in 2010), were caught using mist nets, ringed, morphometric measurements taken, plumage reflectance spectra quantified in the field and feather samples collected for subsequent assessment in the lab.

Colour was quantified based on Endler and Mielke’s [19] approach, using a USB2000 spectrophotometer (Ocean Optics, Dunedin, Florida), and a xenon flash lamp (Ocean Optics). Before using the spectrophotometer, we calibrated it by setting the white and black references, i.e., we “told” the machine which colour we want to be considered as the 100% reflectance (white) standard, and the 0% reflectance (dark) standard, so that the rest of the colour measurements are determined in relation to those maximum and minimum possible reflectance values. We used a WS-1 SS Diffuse Reflectance Standard, a diffuse white plastic >98% reflective from 250-1500 nm, as the white reference (100% reflectance), and a piece of black velvet as the dark standard (0% reflectance) to correct for the noise when no light is reaching the sensor. At the far end of the reflection probe/light source, we put a non-reflective black pointer cut in a 45 degree angle, to avoid mismeasurement derived from the white light directly reflected by the plumage reaching the sensor [15]. Using the spectra acquisition software package OOIBase (Ocean Optics), we measured the reflectance of four body regions, namely the throat, breast, belly and vent of each bird. For the measurements of feather samples in the lab, we collected enough feathers from live birds as to being able to mount them one on top of the other and simulate the original pattern found on live birds. We mounted the feathers on a piece of black velvet to avoid background noise. Once we had tested for the reliability of the measures obtained with both sampling methods separately, we averaged the three measurements for each method and used these average values to test the comparability between field and lab measurements.

We also used Endler and Mielke’s [19] method to calculate brightness, chroma and hue, parameters generally used to specify a colour. Using their equations and the mathematical software Matlab (The MathWorks Inc., Natick, MA), we got the spectral sensitivity functions of the cones corrected for the cut points of oil droplets, calculated the quantal catch for each photoreceptor and converted those quantal catches into dimensional colour space coordinates in a tetrahedral colour space (Fig. 1). Chroma is defined as the strength of the colour signal or the degree of difference in stimulation among the cones, and it is proportional to the Euclidean distance from the origin, that is, the distance from the bird grey (achromatic) point to each point, specified by three space coordinates. Perception of hue depends of which cones are stimulated, and in tetrahedral colour space, it is defined by the angle that a point makes with the origin. As bird colour space is 3D, hue is defined by two angles, analogous to latitude and longitude in geography [19].

**Figure 1:**
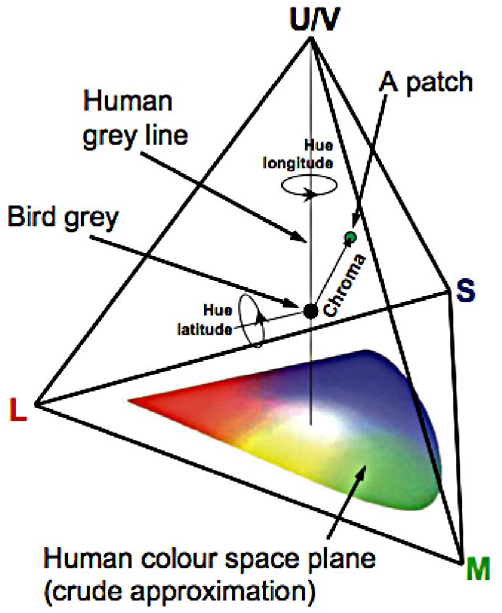
The avian tetrahedral colour space (from Endler and Mielke, 2005).

Brightness is defined as the summed mean reflectance across the entire spectral range (R_300–700_; [44,45]). As well as these parameters, we included UV chroma, a measure of spectral purity, into our analysis, which was calculated as the proportion of reflectance in the UV part of the spectrum (R_300-400_) in relation to the total reflectance spectrum (R_300–700_; [46]). Cone sensitivities and oil droplet cut points were taken from Bowmaker et al. [16], Hart [47], Vorobyev et al. [48], Govardovskii et al. [49], and Hart and Vorobyev [50].

Although all the avian families investigated show plumages reflecting significant amounts of UV light (see [51] for a review), in the particular case of barn swallows, ventral plumage shows a noisy reflectance pattern in the UV part of the spectrum and does not exhibit a clear ultraviolet reflectance peak (Fig. 2; [34]). For this reason, we calculated the same colour variables both including and not including the UV part of the reflectance spectrum, and carried out a repeatability assessment using either the whole reflectance spectrum (300-700 nm) or only the human-visible part (400-700 nm). When using only the visible part, we did not include UV chroma, for obvious reasons, nor hue, as values are identical in both cases.

**Figure 2:**
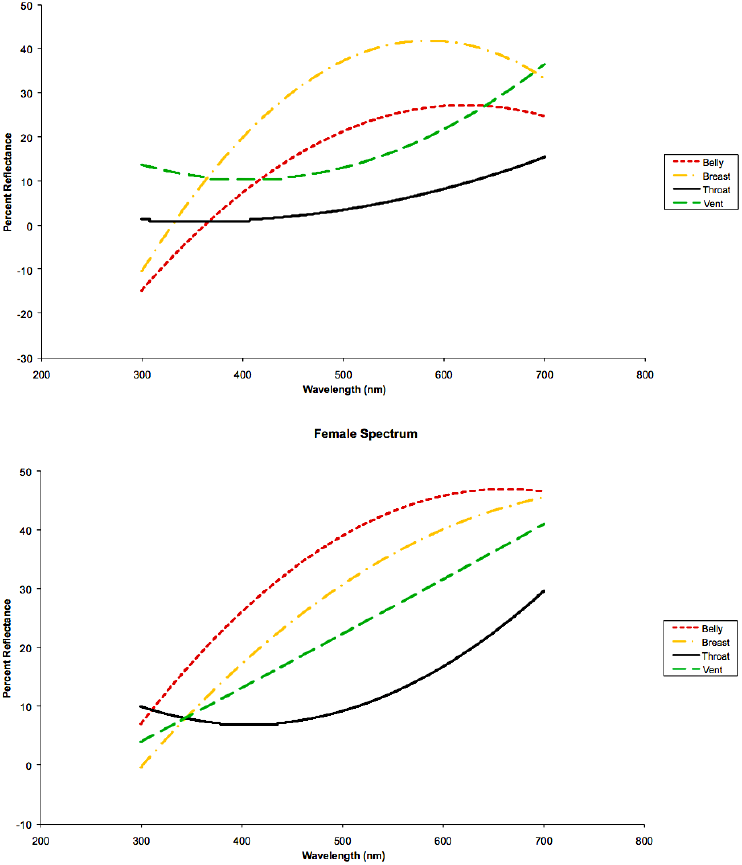
Reflectance spectra for belly, breast, throat and vent patches of male and female barn swallow (regression lines).

Due to the way data were collected, the three plumage colouration measurements taken in the field in 2009 covered a wider area of each plumage patch than the measurements made on feather samples, which were restricted to the area covered by the bunch of feathers plucked from each patch on each individual. In 2010, however, the three field measurements were taken approximately in the same plumage area for each patch.

### ANOVA-based method

In order to test for the reliability of both sampling procedures (described above) separately, we calculated the repeatability for colour variables in the four patches for the different procedures according to Lessells and Boag [43], Senar [52] and Quesada and Senar [39]. Repeatabilities were computed from the mean squares of ANOVA on three repeated measures per individual. Both in field and lab procedures, the second and third measurements were made after removing the reflection probe/light source and the black pointer on top of it and placing it again on the colouration patch. IV took all the measurements.

Once we calculated the “within-method” repeatabilities, i.e., the repeatabilities for each sampling method, we averaged the three measurements per patch per individual and assessed the “between-method” repeatability, i.e. the repeatability of measurements across procedures, but this time the ANOVA was carried out on two repeated measures per individual, one from the field and another one from the lab. We repeated this process for both 2009 and 2010 data separately.

### GLMM-based method

We used a modified version of the R function *R.Anson*, which is itself a modification of rpt.remlLMM function [41]. We fitted two random-effect terms (individual identity and year) in our linear mixed-effects models, and calculated the adjusted repeatability estimate as:

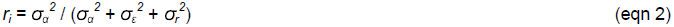

where *σ_r_^2^* is the year variance.

In order to have a general idea of repeatability for each patch, we included all the colour variables in a principal component analysis (PCA) and calculated the repeatability (and confidence intervals) for the first component (PC1) within each plumage patch.

As we conducted multiple statistical tests on data subsets that are not likely to be biologically independent of each other (i.e. different components of the spectra, same metrics on different years, or same metrics in the lab and in the field), there was an increased probability of type I error rates. To control for this increased probability, we corrected our p-values for multiple testing based on the sequentially rejective Bonferroni procedure of Holm [53], using the *p.adjust* function from the *stats* package in R [54]. All the statistical analyses were carried out using R [54,55].

### Ethics Statement

AMcG and MRE had a Home licence which covered taking feather samples as well as other activities (Home Office Project Licence Number 30/2740). MRE was the project licence holder.

All work was carried out on private residencies and farms with the express permission of the landowners in question. Contact details of the landowners can be provided by the authors upon request and after asking the landowners for permission, in order to respect their privacy.

Our field study did not involve any endangered or protected species. The specific locations of the study are provided as supporting material.

Birds were caught using mist nets under licence (AMcG BTO A licence Holder No.4947).

## Results

### ANOVA analyses

In 2009, when including the whole spectrum in the analyses, measuring plumage colouration in the lab proved to be a reliable method. Brightness, UV chroma, chroma and hue latitude and longitude being highly repeatable for all the patches, and hue latitude in the throat being less repeatable (*r*_i_=0.418, *F*_21,44_=3.157, *P*=0.01; Table 1). The method of measuring the plumage colouration in the field (in three different points, covering a wider area of each patch) was also quite consistent but with overall lower values of repeatability, although still reasonably high, for all the variables and patches, being especially low for hue latitude in breast (*r*_i_=0.382, *F*_20,42_=2.856, *P*=0.025) and vent (*r*_i_=0.394, *F*_21,44_=2.955, *P*=0.017; Table 1).

**Table 1:** ANOVA-derived Repeatabilities in 2009 plumage colouration measurements taken from live birds in the field, feather samples in the lab, and across both procedures (UV+Visible spectrum).

|  | UV+Visible range |  |  |  |  |  |  |  |
| --- | --- | --- | --- | --- | --- | --- | --- | --- |
|  | Belly |  | Breast |  | Throat |  | Vent |  |
|  | *F* | *r<sub>i</sub>* | *F* | *r<sub>i</sub>* | *F* | *r<sub>i</sub>* | *F* | *r<sub>i</sub>* |
| **Repeatability field** |  |  |  |  |  |  |  |  |
| Brightness | 6.0446 | **0.627\*\*\*** | 3.1183 | **0.414\*** | 12.273 | **0.79\*\*\*** | 6.1868 | **0.634\*\*\*** |
| UV Chroma | 10.203 | **0.754\*\*\*** | 14.171 | **0.814\*\*\*** | 13.861 | **0.811\*\*\*** | 5.3196 | **0.59\*\*\*** |
| Chroma | 9.0161 | **0.728\*\*\*** | 11.238 | **0.773\*\*\*** | 10.714 | **0.764\*\*\*** | 4.5611 | **0.543\*\*\*** |
| Hue latitude | 4.0921 | **0.508\*\*\*** | 2.8564 | **0.382\*** | 4.4146 | **0.532\*\*\*** | 2.9545 | **0.394\*** |
| Hue longitude | 13.025 | **0.8\*\*\*** | 9.1422 | **0.731\*\*\*** | 24.348 | **0.886\*\*\*** | 4.6771 | **0.55\*\*\*** |
| **Repeatability lab** |  |  |  |  |  |  |  |  |
| Brightness | 13.188 | **0.802\*\*\*** | 14.777 | **0.821\*\*\*** | 8.2092 | **0.706\*\*\*** | 20.212 | **0.865\*\*\*** |
| UV Chroma | 46.493 | **0.938\*\*\*** | 23.015 | **0.88\*\*\*** | 34.895 | **0.919\*\*\*** | 25.561 | **0.892\*\*\*** |
| Chroma | 42.489 | **0.932\*\*\*** | 26.975 | **0.896\*\*\*** | 62.481 | **0.954\*\*\*** | 29.052 | **0.903\*\*\*** |
| Hue latitude | 11.986 | **0.785\*\*\*** | 6.5101 | **0.647\*\*\*** | 3.1573 | **0.418\*\*** | 23.024 | **0.88\*\*\*** |
| Hue longitude | 25.986 | **0.893\*\*\*** | 9.2833 | **0.734\*\*\*** | 6.0119 | **0.625\*\*\*** | 27.291 | **0.898\*\*\*** |
| **Comparison field-lab** |  |  |  |  |  |  |  |  |
| Brightness | 1.4636 | **0.188** | 2.0702 | **0.349** | 0.865 | **-0.072** | 1.8043 | **0.287** |
| UV Chroma | 1.3153 | **0.136** | 1.3591 | **0.152** | 0.7499 | **-0.143** | 0.9006 | **-0.052** |
| Chroma | 0.327 | **-0.507** | 1.3715 | **0.157** | 1.0143 | **0.007** | 0.1376 | **-0.758** |
| Hue latitude | 0.9084 | **-0.048** | 0.8635 | **-0.073** | 1.2523 | **0.112** | 0.706 | **-0.172** |
| Hue longitude | 1.7364 | **0.269** | 1.7513 | **0.273** | 0.6709 | **-0.197** | 1.1055 | **0.05** |

\*\*\*\* $P < 0.001$ ; \*\*\* $P < 0.01$ ; \*\* $P < 0.05$ ; \* $P < 0.1$

When including only the visible part of the spectrum in the analysis, overall repeatability declined. The lab method again proved to be the most reliable, with high values of repeatability for brightness and chroma in all the patches. The field procedure was moderately repeatable for belly and throat, but showed low repeatability for brightness in the breast (*r*_i_=0.36, *F*_20,42_=2.6885, *P*=0.038) and for chroma in the vent (*r*_i_=0.428, *F*_21,44_=3.241, *P*=0.007; Table 2).

**Table 2:** ANOVA-derived Repeatabilities in 2009 plumage colouration measurements taken from live birds in the field, feather samples in the lab, and across both procedures (Only Visible part of the spectrum).

|  | Visible range |  |  |  |  |  |  |  |
| --- | --- | --- | --- | --- | --- | --- | --- | --- |
|  | Belly |  | Breast |  | Throat |  | Vent |  |
| | F | $r_i$ | F | $r_i$ | F | $r_i$ | F | $r_i$ |
| **Repeatability field** |  |  |  |  |  |  |  |  |
| Brightness | 4.8958 | **0.565\*\*\*** | 2.6885 | **0.36\*** | 11.221 | **0.773\*\*\*** | 5.6072 | **0.606\*\*\*** |
| Chroma | 6.7724 | **0.658\*\*\*** | 6.4868 | **0.646\*\*\*** | 6.6207 | **0.652\*\*\*** | 3.2416 | **0.428\*\*** |
| **Repeatability lab** |  |  |  |  |  |  |  |  |
| Brightness | 11.818 | **0.783\*\*\*** | 15.892 | **0.832\*\*\*** | 6.9936 | **0.666\*\*\*** | 19.326 | **0.859\*\*\*** |
| Chroma | 28.865 | **0.903\*\*\*** | 21.117 | **0.821\*\*\*** | 29.402 | **0.873\*\*\*** | 23.838 | **0.901\*\*\*** |
| **Comparison field-lab** |  |  |  |  |  |  |  |  |
| Brightness | 1.461 | **0.187** | 2.0097 | **0.335** | 0.6804 | **-0.19** | 1.8902 | **0.308** |
| Chroma | 2.0563 | **0.346** | 1.6683 | **0.25** | 1.1396 | **0.065** | 1.8415 | **0.296** |

\*\*\*\*) $P < 0.001$ ; \*\*\*) $P < 0.01$ ; \*\*) $P < 0.05$ ; \*) $P < 0.1$ .

The values of repeatability across the field and laboratory procedures were very low for all the patches measured, both considering the whole reflectance spectrum or only the visible part (r_i_<0.35 and *P*>0.05 in all cases), suggesting a lack of consistency across the two assessment methods for melanin-based plumage colouration. Repeatabilities of brightness measurements were slightly higher for the whole spectrum than for only the visible part, in all the patches except from the vent. However, in this case, including only the visible part of the spectrum yielded much more repeatable chroma values than including the whole range, sometimes even turning negative repeatability values into positive, e.g. for the belly (whole range: *r*_i_=-0.507, *F*_20,21_=0.327, *P*=1; only visible range: *r*_i_=0.346, *F*_20,21_=2.056, *P*=0.599), or the vent (whole range: *r*_i_=-0.758, *F*_21,22_=0.138, *P*=1; only visible range: *r*_i_=0.296, *F*_21,22_=1.842, *P*=0.657; Table 1 and Table 2).

In 2010, when including the whole spectrum in the analyses, repeatability measurements in the field (taken approximately in the same point within each patch) yielded considerably higher results than in 2009, with all the r_i_ values above 0.60, except from hue latitude in the throat (r_i_=0.515, F_37,75_=4.186, *P*<0.0001), and with most of the values ranging from 0.74 to 0.91, except for brightness in the breast (r_i_=0.611, F_37,76_=5.710, *P*<0.0001), hue latitude in the belly (r_i_=0.63, F_37,76_=6.1, *P*<0.0001), hue latitude in the vent (r_i_=0.629, F_37,76_=6.088, *P*<0.0001) and hue longitude in the vent (r_i_=0.679, F_37,76_=7.356, *P*<0.0001). In the lab, repeatability values ranged from 0.71 to 0.95 in most of the patches, except from hue latitude in the throat (r_i_=0.65, F_37,76_=6.569, *P*<0.0001), and repeatability was overall higher than when measuring it on live birds, except from UV chroma in the belly (r_i_=0.857, F_37,76_=18.986, *P*<0.0001), breast (r_i_=0.788, F_37,76_=12.117, *P*<0.0001) and vent (r_i_=0.819, F_37,76_=14.571, *P*<0.0001) and hue latitude in the breast (r_i_=0.722, F_37,76_=8.808, *P*<0.0001), where it was slightly lower. Repeatability values were similar to the ones obtained in 2009 (Table 3).

**Table 3:** ANOVA-derived repeatabilities in 2010 plumage colouration measurements taken from live birds in the field, feather samples in the lab, and across both procedures (UV+Visible spectrum).

|  | UV+Visible range |  |  |  |  |  |  |  |
| --- | --- | --- | --- | --- | --- | --- | --- | --- |
|  | Belly |  | Breast |  | Throat |  | Vent |  |
|  | F | r<sub>i</sub> | F | r<sub>i</sub> | F | r<sub>i</sub> | F | r<sub>i</sub> |
| **Repeatability field** |  |  |  |  |  |  |  |  |
| Brightness | 16.422 | **0.837\*\*\*** | 5.7104 | **0.611\*\*\*** | 17.398 | **0.846\*\*\*** | 14.534 | **0.819\*\*\*** |
| UV Chroma | 25.333 | **0.89\*\*\*** | 15.853 | **0.832\*\*\*** | 12.357 | **0.791\*\*\*** | 16.829 | **0.841\*\*\*** |
| Chroma | 24.037 | **0.885\*\*\*** | 14.749 | **0.821\*\*\*** | 22.212 | **0.876\*\*\*** | 12.377 | **0.792\*\*\*** |
| Hue latitude | 6.0997 | **0.630\*\*\*** | 9.7411 | **0.744\*\*\*** | 4.1857 | **0.515\*\*\*** | 6.0878 | **0.629\*\*\*** |
| Hue longitude | 23.669 | **0.883\*\*\*** | 31.363 | **0.910\*\*\*** | 9.8914 | **0.748\*\*\*** | 7.3556 | **0.679\*\*\*** |
| **Repeatability lab** |  |  |  |  |  |  |  |  |
| Brightness | 55.197 | **0.947\*\*\*** | 31.387 | **0.910\*\*\*** | 30.9 | **0.909\*\*\*** | 44.036 | **0.934\*\*\*** |
| UV Chroma | 18.986 | **0.857\*\*\*** | 12.117 | **0.788\*\*\*** | 17.544 | **0.847\*\*\*** | 14.571 | **0.819\*\*\*** |
| Chroma | 25.375 | **0.890\*\*\*** | 25.936 | **0.893\*\*\*** | 27.679 | **0.899\*\*\*** | 39.854 | **0.928\*\*\*** |
| Hue latitude | 8.3908 | **0.711\*\*\*** | 8.8076 | **0.722\*\*\*** | 6.5692 | **0.650\*\*\*** | 9.1422 | **0.731\*\*\*** |
| Hue longitude | 37.357 | **0.924\*\*\*** | 31.683 | **0.911\*\*\*** | 11.664 | **0.781\*\*\*** | 37.517 | **0.924\*\*\*** |
| **Comparison field-lab** |  |  |  |  |  |  |  |  |
| Brightness | 1.3041 | **0.131948** | 2.8612 | **0.482\*** | 0.7642 | **-0.134** | 2.0703 | **0.349** |
| UV Chroma | 0.5572 | **-0.284** | 1.059 | **0.029** | 1.9441 | **0.321** | 1.5873 | **0.227** |
| Chroma | 0.9059 | **-0.049** | 1.9662 | **0.326** | 1.755 | **0.274** | 1.5788 | **0.224** |
| Hue latitude | 2.5152 | **0.431\*** | 1.2479 | **0.110243** | 0.7839 | **-0.121** | 2.7232 | **0.463\*** |
| Hue longitude | 8.7322 | **0.794\*\*\*** | 4.8175 | **0.657\*\*\*** | **0.9731** | **-0.014** | 3.5525 | **0.561\*\*** |

\*\*\*\*) $P < 0.001$ ; \*\*\*) $P < 0.01$ ; \*) $P < 0.05$ ; .) $P < 0.1$

When doing the analysis including only the visible part of the spectrum, measuring colouration in the lab was again the most reliable method of both, with all the r_i_ values above 0.91, except for chroma in the throat (r_i_=0.881, F_37,76_=22.964, *P*<0.0001). Field procedure still yielded high repeatability values, with brightness in the breast (r_i_=0.569, F_37,76_=4.959, *P*<0.0001) and chroma in the vent (r_i_=0.696, F_37,76_=7.839, *P*<0.0001) being the only measurements with values below 0.81. For both methods, repeatability values were higher than in 2009 for all the variables within all the plumage patches (Table 4).

**Table 4:** ANOVA-derived Repeatabilities in 2010 plumage colouration measurements taken from live birds in the field, feather samples in the lab, and across both procedures (Only Visible part of the spectrum).

|  | Visible range |  |  |  |  |  |  |  |
| --- | --- | --- | --- | --- | --- | --- | --- | --- |
|  | Belly |  | Breast |  | Throat |  | Vent |  |
|  | F | r<sub>i</sub> | F | r<sub>i</sub> | F | r<sub>i</sub> | F | r<sub>i</sub> |
| **Repeatability field** |  |  |  |  |  |  |  |  |
| Brightness | 14.408 | **0.817\*\*\*** | 4.959 | **0.569\*\*\*** | 16.59 | **0.839\*\*\*** | 13.527 | **0.806\*\*\*** |
| Chroma | 22.566 | **0.878\*\*\*** | 22.163 | **0.876\*\*\*** | 20.007 | **0.864\*\*\*** | 7.8386 | **0.696\*\*\*** |
| **Repeatability lab** |  |  |  |  |  |  |  |  |
| Brightness | 59.713 | **0.951\*\*\*** | 34.778 | **0.918\*\*\*** | 34.133 | **0.917\*\*\*** | 49.55 | **0.942\*\*\*** |
| Chroma | 51.455 | **0.944\*\*\*** | 57.897 | **0.950\*\*\*** | 22.964 | **0.881\*\*\*** | 72.314 | **0.959\*\*\*** |
| **Comparison field-lab** |  |  |  |  |  |  |  |  |
| Brightness | 1.5559 | **0.217** | 3.1819 | **0.522\*\*** | 0.679 | **-0.191** | 2.3696 | **0.406.** |
| Chroma | 7.0047 | **0.750\*\*\*** | 7.0261 | **0.751\*\*\*** | 1.0876 | **0.042** | 2.4275 | **0.416\*** |

\*\*\*\*) $P < 0.001$ ; \*\*\*) $P < 0.01$ ; \*\*) $P < 0.05$ ; \*) $P < 0.1$

Repeatabilities across field and lab methods in 2010 were quite heterogeneous including the whole spectrum in the analyses: moderately high for hue longitude in the belly (r_i_=0.794, F_37,38_=8.732, *P*<0.0001) and breast (r_i_=0.657, F_37,38_=4.818, *P*<0.0001), moderate for vent hue latitude (r_i_=0.463, F_37,38_=2.723, *P*=0.018) and longitude (r_i_=0.561, F_37,38_=3.553, *P*=0.001), belly hue latitude (r_i_=0.431, F_37,38_=2.515, *P*=0.034) and breast brightness (r_i_=0.482, F_37,38_=2.861, *P*=0.012), and low for breast chroma (r_i_=0.326, F_37,38_=1.966, *P*=0.21), throat UV chroma (r_i_=0.321, F_37,38_=1.944, *P*=0.312) and chroma (r_i_=0.274, F_37,38_=1.755, *P*=0.576) and vent brightness (r_i_=0.349, F_37,38_=2.070, *P*=0.141). For the rest of the cases, repeatabilities were very low (r_i_<0.23 and *P*>0.05 for all the cases). When including only the visible part of the spectrum, repeatability was moderate to high and had a significant effect for both brightness and chroma in the breast, and for chroma in the vent and in the belly, whereas it was quite low and non-significant for brightness in the belly (r_i_=0.217, F_37,38_=1.556, *P*=0.899), and very low for both variables in the throat. Repeatability values for chroma in all the patches except for the throat were much higher than when we included the whole spectrum range, e.g. in the belly (whole range: r_i_=-0.049., F_37,38_=0.906, *P*=1; only visible range: r_i_=0.75, F_37,38_=7.005, *P*<0.0001) and in the vent (whole range: r_i_=0.224, F_37,38_=1.579, *P*=0.657; only visible range: r_i_=0.416, F_37,38_=2.428, *P*=0.047; Table 3 and Table 4).

### Generalized Linear Mixed Model analyses

When we included the whole spectrum in the analyses, for all the principal component analyses carried out within each patch for field and lab measurements, PC1 accounted for more than a 53% of the total variance, except for vent measurements in the field (where it explained a 49% of the total variance) and for measurements in the throat (where it explained between 44% and 47%). When including only the visible part of the spectrum, PC1 explained a 67% of the total variance for vent measurements in the field, and between 75% and 93% in the rest of the cases.

Repeatability of feather colour measurements was much higher and confidence intervals smaller when quantifying colouration from measurements taken in the lab than when doing it on measurements taken on live birds in the field, both including the whole spectrum in the analyses or only the visible part, being particularly high in the breast (whole range: r_i_=0.916, 95%CI=0.855, 0.943, *P*<0.0001; visible range: r_i_=0.927, 95%CI=0.872, 0.95, *P*<0.0001). All the repeatability values from lab measurements ranged between 0.71 and 0.93 and were highly significant (*P*<0.0001).

When using field measurements, repeatabilities were still moderately high (all r_i_ values above 0.50) and higher when including only the visible spectrum range in the analyses than including the whole range, except in the throat (whole range: r_i_=0.629, 95%CI=0.303, 0.874, *P*<0.0001; visible range: r_i_=0.564, 95%CI=0.266, 0.816, *P*<0.0001; Fig. 3).

**Figure 3:**
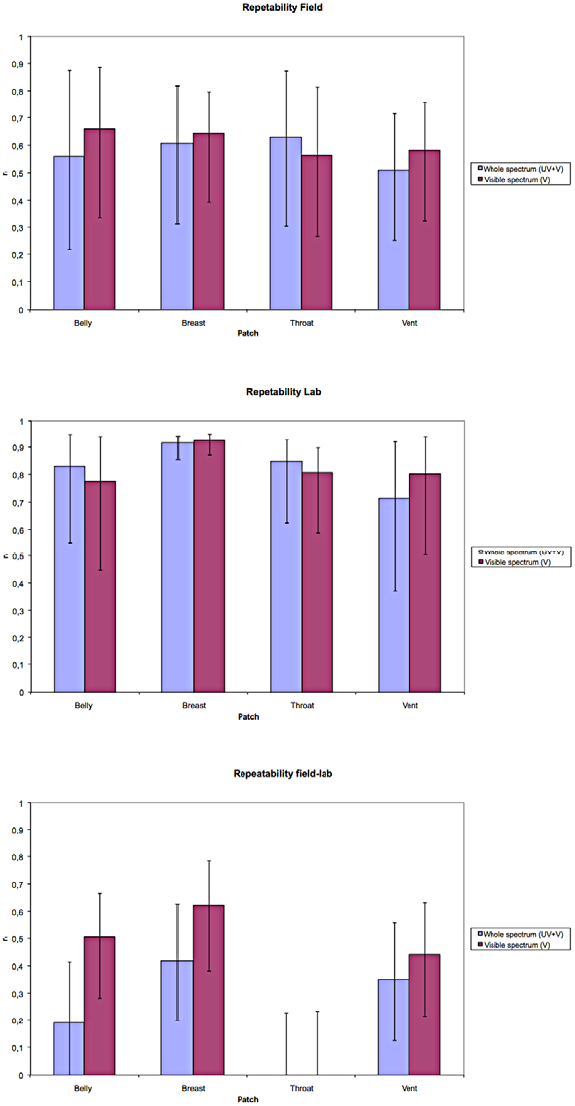
GLMM-derived repeatabilities (± 95% CI) in 2009-2010 plumage colouration measurements taken a) from live birds in the field, b) from feather samples in the lab, and c) across both procedures, both when including the whole light spectrum or only the human-visible spectrum in the analyses.

Repeatabilities across both field and lab methods yielded higher results when we included only the visible spectrum in the analyses than when we included the whole spectrum. Leaving the values for throat apart, as here repeatability was not significantly different from zero no matter the spectrum range we included in the analyses, r_i_ ranged between 0.19 and 0.41 when including the whole spectrum, and between 0.44 and 0.62 when including only the visible part (Fig. 3). All the repeatabilities in belly, breast and vent were significant except for the belly when including the whole spectrum (r_i_=0.189, 95%CI=0, 0.415, *P*=0.069), but it became significant and higher when only the visible spectrum was included in the analyses (r_i_=0.503, 95%CI=0.281, 0.667, *P*<0.0001, Fig. 3).

## Discussion

### Measuring plumage ornamentation to gain repeatable and reliable measures

Measuring plumage colouration from feather samples in the lab proved to be a highly reliable method, with high values of repeatability in general for all the variables and patches in 2009, 2010 and when applying the GLMM-based method for both years. Measurements taken directly on specimens in the field showed a reasonable extent of reliability, but with overall lower values of repeatability compared to the lab for most of the variables measured on different patches. As an exception, some variables in the throat in 2009 and UV chroma measurements in the belly, breast and vent in 2010 when considering the whole spectrum, yielded higher values of repeatability when measured in the field, only when applying the ANOVA-based method.

A potential explanation for our findings is that the throat patch is smaller and much darker than the rest of the patches. The feathers of the throat patch are also considerably smaller. Therefore, it is often quite difficult to obtain a reliable reflectance measurement with such a limited amount of photons reaching the spectrophotometer probe. Also, it is more difficult to create a “plumage patch” in the lab with a feather arrangement similar to the bird’s original one and big enough to be able to apply the spectrophotometer pointer to it. An alternative explanation is that the UV part of the spectrum shows a highly noisy pattern, thus highly consistent UV chroma repeatabilities across field or lab measurements may not necessarily be expected. This latter alternative may explain why repeatability values in the lab for UV chroma measurements were higher in the field in 2010, whereas the rest of the repeatability values consistently tended to be higher in the lab.

When applying the GLMM-based method, repeatability was moderate to high within all the patches for field measurements, whereas it was considerably higher, and the confidence intervals considerably narrower, for lab measurements. This finding suggests that lab-based measures are more reliable ways of assessing melanin-based colouration.

When comparing both methods, the values obtained in 2009 for different variables measured in different plumage patches directly on field birds were poorly repeatable compared to the values obtained for the same variables measured from feather samples in the lab, and non-significant in all cases. In 2010, in contrast, repeatabilities were higher and significant for certain metrics in certain patches only. These results stand in marked contrast to the positive results of another study, which compared the repeatabilities between both colouration assessment procedures for carotenoid-based plumage [39]. There can be several reasons for this difference: for example, due to the different characteristics of the two types of pigments, carotenoid-derived colouration is more variable among individuals than melanin-based colouration [56], and repeatability of a character increases with variability [52]. In order to increase the repeatability of some measurements, a possible solution could be to increase the number of measurements, for example from three to five, as it has already been done by several authors [22,28,32]. Unfortunately, this is not an option when working with live birds in the field, as we would be increasing the manipulation times and, consequently, the stress levels to an unacceptable degree, although it can be applied when assessing colouration in the lab on feather samples [39].

As mentioned above, the three plumage colouration measurements taken in the field in 2009 covered a wider area of each plumage patch than in 2010, when the three field measurements were taken approximately in the same plumage area for each patch, being this area also the one from which the majority of feathers for lab measurements were taken. Due to the different ways in which data were collected in the field each year, repeatability of 2009 field measurements can be taken as an estimate of colouration consistency within the plumage patches. Our results suggest a moderate to high within-patch consistency for melanin-based ventral colouration in our model species. The comparability of both procedures in 2009 may have been compromised, although the repeatability of the 2010 samples was higher even for lab measurements, especially when considering only the visible part of the spectrum. This may be indicative of higher patch colouration homogeneity in 2010. Also, the higher lab repeatabilities in 2009 than the field ones could be argued to be a function of field measures being taken over bigger plumage areas. But the fact that we still get higher lab than field measurements in 2010 suggests practical limitations to repeatability measurements in the field.

Consequently, collecting feathers from birds and assessing their colouration in the lab, as well as being more convenient, minimising risk to a sensitive device like a spectrophotometer and reducing handling times of the animals [39], is a more reliable method for assessing melanin plumage colouration than doing so directly on live birds, according to our results.

### Assessing repeatability

The GLMM-based method [41], applied to data from both years, allowed us to control for year effects by adding the year variance into the total variance calculation, so that we could obtain the adjusted repeatability for data from both years. The PCA allowed to create composite variables accounting for almost 50% of the total variance in the metrics taken from each patch, which therefore, made it possible to estimate the overall repeatability within each patch for both methods separately and across methods.

The possibility of calculating adjusted repeatabilities by including year as a random factor, together with the reduction in the number of variables accounting for a great proportion of the total variance achieved by the PCA, considerably reduced the amount of multiple tests necessary for repeatability calculation. Thus, the p-values obtained with this method were less affected by Bonferroni corrections than those obtained with the ANOVA-based method, reducing the probability of type II errors and increasing the power of this repeatability-calculation method.

Additionally, this reduction in the number of variables offered us the possibility of getting a general view of the colour measurement repeatability within each patch. As our variables are all derived from the reflectance spectra obtained on each colouration patch through the spectrophotometer, comparing the repeatability of all the different variables within the patches, as well as increasing the probability of type II errors, can be redundant. The PCA-GLMM combination, in contrast, made it possible to compare the first principal components within each patch, which can be regarded as whole spectrum estimates, as they gather a great proportion of the total variance contained in all the variables originally extracted from the reflectance spectra measurements. And it also allowed us to do so with data from both years.

Finally, thanks to the use of the GLMM-based method, we could calculate confidence intervals, useful indicators of the reliability of our repeatability estimates, additionally to just p-values. That way, and together with the advantages mentioned above, it was possible to get a more complete and reliable overall perspective of the question being studied.

The fact that almost all the repeatability measurements, and especially the repeatabilities across field and lab methods (in patches other than the throat), were higher when including only the human-visible spectrum in the analyses, suggests that the noisy reflectance pattern in the UV part of the spectrum may be decreasing the repeatability and underestimating the comparability of the two methods. For throat plumage, however, we observed the opposite trend, with higher repeatability values when including the whole spectrum, which could be indicative that the UV part of the spectrum is more important in the throat than in the rest of the patches. We find support for this idea when looking at reflectance spectra plots for different patches (Fig. 2): throat reflectance spectra, although showing also quite a noisy pattern for the UV part, and unlike the rest of the patches’ spectra, tends to show UV reflectance peaks in both sexes. Further work is needed to find out whether there are UV reflectance differences amongst different plumage patches.

## Conclusions

Our results suggest that collecting feathers from live animals and assessing colouration in the lab is a better approach for measuring plumage ornamentation in order to gain repeatable and reliable results compared to direct measures on live birds in the field. In addition, since it is easier on equipment and minimises the length of time birds need to be handled (minimising the stress levels inflicted on them), feather sampling would appear to be the best method available.

In addition, from a statistical point of view, our results support the superiority of the GLMM-based method [41] for repeatability calculation, as it enables random factors to be accounted for and can calculate adjusted repeatability values, which are more accurate than those calculated using other (e.g., ANOVA) methods and increase the power of the tests. The reduction in the number of variables gives us a general, patch by patch overview of the problem being studied, and the confidence intervals allow us to test the reliability of our own repeatability estimates.

Finally, we have also shown that it is important to check for the effect that the UV part of the spectrum could be exerting on repeatability calculations, as the capability of the plumage to reflect the UV light could have different biological implications in different plumage patches.

## Acknowledgements

Thanks to Mary-Anne Collis, who contributed to establish the farm network, and Ian Blessley, who extended the network and initially talked to some of the farmers. Our gratitude to Shinichi Nakagawa, who helped us with the modification of his R function for our purposes. Rebecca Safran was a huge inspiration for our work. Jon Blount introduced I.V.A. to the use of the spectrophotometer, and Tom Pike allowed us to use his Matlab spectral data processing equations. Thanks to Joan Carles Senar for his invaluable help with repeatability-related issues. We are very much indebted to all the farmers and landlords who granted us access to their properties, their lovely families, neighbours and pets. Two anonymous reviewers provided helpful comments on a previous version of the manuscript.

